# Obesity promotes Fumonisin B1 toxicity and induces hepatitis

**DOI:** 10.1101/2022.07.22.500801

**Authors:** Léonie Dopavogui, Marion Régnier, Arnaud Polizzi, Quentin Ponchon, Sarra Smati, Wendy Klement, Frédéric Lasserre, Céline Lukowicz, Yannick Lippi, Anne Fougerat, Justine Bertrand-Michel, Claire Naylies, Cécile Canlet, Laurent Debrauwer, Laurence Gamet-Payrastre, Charlène Dauriat, Josefina Casas, Siska Croubels, Siegrid De Baere, Hester M. Burger, Benoit Chassaing, Sandrine Ellero-Simatos, Hervé Guillou, Isabelle P. Oswald, Nicolas Loiseau

## Abstract

**Background and aim:** Obesity is a major public health issue worldwide. Obesity is associated with chronic inflammation that contribute to long-term complications, including insulin resistance, type 2 diabetes and non-alcoholic fatty liver disease. We hypothesized that obesity may also influence the sensitivity to food contaminants, such as fumonisin B1 (FB1), a mycotoxin produced mainly by the *Fusarium verticillioides*. FB1, a common contaminant of corn, is the most abundant and best characterized member of the fumonisins family. This toxin provokes severe mycotoxicosis in animals, which leads to hepatotoxicity and alterations in the immune response and intestinal barrier permeability. We investigated here whether diet-induced obesity could modulate the sensitivity to oral FB1 exposure, with emphasis on gut health and hepatotoxicity.

**Methods:** The metabolic effects of FB1 were assessed in obese and non-obese male C57BL/6J mice. For 15 weeks, mice received a high-fat diet (HFD) or normal chow diet (CHOW). During the last three weeks, mice were exposed or not to FB1 (10 mg/kg body weight/day) through drinking water.

**Results:** As expected, HFD feeding induced significant body weight gain, glucose intolerance, and hepatic steatosis. FB1-exposed mice displayed a higher sphinganine/sphingosine ratio, a well-known FB1 biomarker of exposure, due to inhibition of ceramide synthases activity by FB1. Combined exposure to HFD and FB1 resulted in body weight loss and a decrease in fasting blood glucose level. This co-exposition also induces gut dysbiosis, an increase in plasma FB1 level, a decrease in liver weight and hepatic steatosis. Moreover, plasma transaminase levels were significantly increased and associated with liver inflammation in HFD/FB1-treated mice. Liver gene expression analysis revealed that the combined exposure to HFD and FB1 was associated with reduced expression of genes involved in lipogenesis and increased expression of immune response and cell cycle-associated genes.

**Conclusion:** These results suggest that, in the context of obesity, FB1 exposure promotes gut dysbiosis and severe liver inflammation. To our knowledge, this study provides the first example of obesity-induced hepatitis in response to a food contaminant.

## 1. Introduction

The prevalence of obesity has reached 13% of the adult population worldwide, and 39% of the world’s adult population is considered overweight (WHO, 2021). Therefore, obesity is considered as an epidemic disease and represents a major public health burden worldwide. Obesity promotes many other diseases, such as type 2 diabetes, cardiovascular diseases, and non-alcoholic fatty liver disease (NAFLD). Obesity fosters disease development through a combination of metabolic changes (Cirulli et al., 2018) and chronic low-grade inflammation (Rohm et al., 2022). Obesity is highly linked to lifestyle and the environment. High-caloric-density diets and reduced physical activities are thought to play an important role in the development of this epidemic. In addition to genetic factors, many environmental factors influence obesity (Pillon et al., 2021), including xenobiotics, endocrine disruptors (Sun et al., 2022), and other food additives (Chassaing et al., 2015; Suez et al., 2014). Although there is increasing evidence that food contaminants can impact the development of obesity, very few studies have investigated the influence of obesity on the sensitivity to food contaminants.

Mycotoxins are fungal toxins that contaminate animal feed and human food worldwide; thus, they cause significant veterinary and public health issues. *Fusarium* spp. is among the most frequent of the fungal genera found in different cereal crops; it causes economic loss and food safety concerns, because it reduces the cereal yield and quality (Cano et al., 2016). Moreover, climate change has led to shifts in temperature and humidity conditions, which favor *Fusarium* dissemination (Nnadi et al., 2021). Fumonisins are the predominant mycotoxins produced by *Fusarium* spp., and fumonisin B1 (FB1) is the most prevalent and the most documented member of this family (Knutsen et al., 2018a). In 2007, the European Union set recommendations and regulations (Commission Recommendation 2006 [Ec] No 576/2006; Commission Regulation 2007 [Ec] No 1126/2007) for the maximum levels of fumonisins (sum of FB1 and FB2) allowed in animal feed (from 5 mg/kg for pig feed to 50 mg/kg for adult ruminant feed) and human foodstuffs (from 0.2 mg/kg for baby foods to 4 mg/kg for unprocessed maize).

FB1 exposure induces severe mycotoxicosis in pigs (Knutsen et al 2018b), with diverse clinical symptoms. The most common symptoms are nephrotoxicity, hepatotoxicity (Terciolo et al., 2019), immunotoxicity (Devriendt et al., 2009; Halloy et al., 2005), and intestinal barrier function disturbances (Bouhet et al., 2006; Loiseau et al., 2007). To date, the known molecular mechanisms underlying FB1 toxicity are mostly related to its inhibitory effect on sphingolipid biosynthesis (Wang et al., 1991; Chen et al., 2021). Indeed, FB1 and sphingoid long-chain bases share similar structural backbone features. The inhibition of ceramide synthase increases free sphinganine levels and reduces the abundance of complex sphingolipids and ceramides (Loiseau et al., 2007). This effect results in elevating the ratio of free sphingoid bases (sphinganine/sphingosine, Sa/So) in several tissues (e.g., liver and intestine), in plasma, and in cell lines (Grenier et al., 2012; Riley et al., 1993).

Previous studies from our group showed that FB1 had a significant influence on lipid metabolism (Régnier et al., 2017; Régnier et al., 2019). Therefore, the current study aimed to investigate the effect of obesity on FB1 toxicity. We fed mice a high-fat diet (HFD) to induce obesity *in vivo*. Next, we investigated the systemic effects through the evolution of the gut microbiota ecology balance and the hepatic responses to FB1 exposure, in both normal-weight and obese mice. Our work showed that obesity enhanced FB1 plasma levels, which strongly impacted mouse metabolism. In obese mice, FB1 exposure reduced glucose intolerance and reduced steatosis, but promoted severe hepatitis.

## 2. Materials and methods

### 2.1 Animals, diet, and exposure to FB1

All experiments were carried out in accordance with the European Guidelines for the Care and Use of Animals for Research Purposes. The animal study protocol was approved by an independent ethics committee (CEEA-86 Toxcométhique) under the authorization number 2016070116429578. The animals were treated humanely with due consideration to the alleviation of distress and discomfort. Mouse housing was controlled for temperature (21-23°C) and light (12 h light/12 h dark). A total of 48 C57BL/6J male mice (6 weeks old) were purchased from Charles Rivers Laboratories (L’Arbresle, France). Mice were allowed two weeks of acclimatization with free access and *ad libitum* water and food, with a standard rodent diet (safe 04 U8220G10R) from SAFE (Augy, France). Then, the mice were randomly divided into four groups of 12 mice each. Two groups (n=24, 4 cages of 6 mice) were fed a chow diet with 10 kcal% fat (CHOW, D12450J, Research Diets) and the other two groups (n=24, 4 cages of 6 mice) were fed a high-fat diet with 60 kcal% fat (HFD, D12492, Research Diets) for 15 weeks. After 12 weeks of feeding, half of the CHOW (n=12, 2 cages of 6 mice) and HFD (n=12, 2 cages of 6 mice) groups were exposed to FB1 (10 mg/kg bw/day) by adjusting every two days the amount of consumed FB1 in the drinking water during 3 weeks in order to maintain a constant level of exposure. Every week, mice were weighed, and water consumption was measured to adjust the quantity of FB1 in the water. Food intake was also monitored. At the end of the experiment, mice were sacrificed to collect plasma and tissue samples.

### 2.2 Blood and tissue sampling

After 15 weeks of feeding, mice were fasted for 6 h, and blood glucose levels were measured from the tail vein with an AccuCheck Performa glucometer (Roche Diagnostics). At the end of the experiment, blood was collected into EDTA-coated tubes (BD Microtainer, K2E tubes) from the submandibular vein. Plasma was isolated by centrifugation (1500 ×*g* for 10 min at 4°C) and stored at −80°C until use for plasma biochemistry. All mice were sacrificed, on the day 104, in the fed state. Following sacrifice by cervical dislocation, liver and caecum were removed, weighed, prepared for histology analysis or snap frozen in liquid nitrogen and stored at −80°C.

### 2.3 Plasma FB1 Analysis

Equal volumes of plasma of 4 individual mice from each group were pooled and 100 µl was used for analysis. Considering this pooling of samples, only 3 FB1 level analysis have been performed per group. Plasma FB1 was analyzed with a validated UPLC-MS/MS (ultra-performance liquid chromatography-tandem mass spectrometry) method previously described (De Baere et al., 2018). The FB1 analytical standard was provided by Fermentek Ltd (Jerusalem, Israel). The limit of quantification was determined at 0.5 ng/ml, using 100 µl of plasma. The limit of detection, corresponding to a signal-to-noise value of 3/1, was 0.09 ng/ml.

### 2.4 Biochemical analyses

We analyzed the following plasma constituents: alanine aminotransferase (ALT), aspartate aminotransferase (AST), alkaline phosphatase (ALP), bilirubin, creatinine, triglycerides, total cholesterol, high density lipoprotein, and low-density lipoprotein cholesterol. All biochemical analyses were performed with a COBASMIRA+ by the Anexplo technical platform team (I2MC, Rangueil, Toulouse).

### 2.5 Lipid extraction and analysis

Liver samples were homogenized in Lysing Matrix D tubes with 1 ml of methanol/5 mM EGTA (2:1 v/v) in a FastPrep machine (MPBiochemicals). Lipids corresponding to an equivalent of 2 mg of tissue were extracted according to Bligh and Dyer, in chloroform/methanol/water (2.5:2.5:2, v/v/v), in the presence of the following internal standards: glyceryl trinonadecanoate, stigmasterol, and cholesteryl heptadecanoate (Sigma, Saint-Quentin-Fallavier, France). Total lipids were suspended in 160 µl ethyl acetate, and the triglycerides, free cholesterol, and cholesterol ester components were analyzed with FID gas-chromatography on a focus Thermo Electron system with a Zebron-1 Phenomenex fused-silica capillary column (5 m, 0.32 mm i.d., 0.50 mm film thickness). The oven temperature was programmed to increase from 200 to 350°C at a rate of 5°C/min, and the carrier gas was hydrogen (0.5 bar). The injector and the detector were at 315°C and 345°C, respectively.

Liver ceramide, sphingomyelin, sphingosine, and sphinganine were extracted, as previously described (Barbacini et al., 2019), with chloroform/water/methanol (2.5:1:5 v/v/v) in the presence of the following internal standards: ceramide d18:1/12:0 (16 ng), sphingomyelin d18:1/12:0 (16 ng), sphingosine 17:0, and sphinganine 17:0 and sphingosine-1-phosphate 17:0. Sphingolipids and internal standards were analyzed by liquid chromatography mass spectrometry (LC-MS) with an Acquity ultra high-performance liquid chromatography (UHPLC) system (Waters, USA) connected to a Time of Flight (LCT Premier XE, Waters, USA) Detector or a triple quadrupole mass spectrometer (Xevo, Waters, USA). The final data were calculated as pmol/mg of protein.

### 2.6 Proton nuclear magnetic resonance (1H-NMR)-based metabolomics

1H NMR spectroscopy was performed on aqueous liver extracts prepared from liver samples (50–75 mg). Briefly, livers were homogenized in chloroform/methanol/NaCl 0.9% (2/1/0.6, v/v/v) containing 0.1% butyl hydroxytoluene. Homogenates were centrifuged at 5,000 ×*g* for 10 min. The supernatant was collected, lyophilized, and reconstituted in 600 μl of D2O that contained 0.25 mM 3-(trimethylsilyl) propionic-(2,2,3,3-d4) acid sodium salt (TSP), as a chemical shift reference at 0 ppm.

All 1H NMR spectra were obtained on a Bruker DRX-600-Avance NMR spectrometer (Bruker) equipped with the AXIOM metabolomics platform (MetaToul). The instrument was operated at 600:13 MHz for 1H resonance frequency. It included an inverse detection 5-mm 1H-13C-15N cryoprobe attached to a cryoplatform (the preamplifier cooling unit).

1H NMR spectra were acquired at 300 K with a standard, one-dimensional noesypr1D pulse sequence with water presaturation and a total spin-echo delay (2 ns) of 100 ms. Data were analyzed by applying an exponential window function with a 0.3-Hz line broadening, prior to Fourier transformation. The resulting spectra were phased, baseline-corrected, and calibrated to TSP (0:00 ppm) manually with Mnova NMR (version 9.0; Mestrelab Research S.L.). The spectra were subsequently imported into MatLab (R2014a; MathWorks, Inc.). All data were analyzed with the use of full-resolution spectra. The region containing the water resonance (4:6– 5:2 ppm) was removed, and the spectra were normalized to the probabilistic quotient (Dieterle et al. 2006) and aligned with a previously published function (Veselkov et al. 2009).

Data were mean-centered and scaled with unit variance scaling, prior to performing orthogonal projection on latent structure-discriminant analysis (O-PLS-DA). The O-PLS derived model was evaluated for accuracy of prediction (Q2Y value) with 10-fold cross-validation. The parameters of the final models are indicated in the figures. Metabolite identifications and discriminations between the groups were performed by calculating the O-PLS-DA correlation coefficients (r2) for each variable, and then, back-scaling into a spectral domain to preserve the shapes of the NMR spectra and the signs of the coefficients (Cloarec et al. 2005). The weights of the variables were color-coded, according to the square of the O-PLS-DA correlation coefficients.

Correlation coefficients extracted from significant models were filtered, and only significant correlations above the threshold defined by Pearson’s critical correlation coefficient (p<0:05; r^2^>0.55; for n=12 per group) were considered significant. For illustration purposes, the areas under the curves of several signals of interest were integrated, and significance was tested with a univariate test.

### 2.7 Histology

Hematoxylin/eosin (H&E) staining was performed on paraformaldehyde-fixed, paraffin-embedded liver tissue sections (3 µm). Sections were visualized with a Leica DFC300 camera. Livers were examined with light microscopy. First, liver sections were screened to determine all the effects present on each section. The histological features were grouped with the steatosis score (evaluated according to Contos *et al*., 2001). Liver sections were evaluated for steatosis and inflammation. The steatosis score was based on the percentage of hepatocytes that contained fat, where Grade 0 = no hepatocytes containing fat in any section; grade 1 = 1% to 25% of hepatocytes; grade 2 = 26% to 50% of hepatocytes; grade 3 = 51% to 75% of hepatocytes; and grade 4 = 76% to 100% of hepatocytes. The inflammation score was the number of inflammatory foci counted in 10 distinct 200× fields for each liver section. Values represented the mean of 10 fields/liver section.

### 2.8 Gene expression studies

Total cellular RNA was extracted with Trizol reagent (Invitrogen). Transcriptome profiles were performed with the Agilent Whole Mouse Genome microarray (4×44K), according to manufacturer instructions. Microarray data and all experimental details are available in the Gene Expression Omnibus Series database (accession number GSE208735; https://www.ncbi.nlm.nih.gov/geo/query/acc.cgi?acc=GSE208735).

Total RNA samples (2 μg) were reverse-transcribed with the high-capacity cDNA reverse transcription kit (Applied Biosystems), then analyzed with real-time quantitative polymerase chain reaction (qPCR). Primers for the Sybr Green assays are presented in Supplementary Table 1. Amplifications were performed on a Stratagene Mx3005P thermocycler (Agilent Technology). qPCR data were normalized to the endogenous level of proteasome 20S subunit beta 6 messenger RNA (mRNA) and analyzed with LinRegPCR software.

### 2.9 Microbiota composition analysis through 16S rRNA gene sequencing

We performed 16S ribosomal RNA (rRNA) gene amplification and sequencing with Illumina MiSeq technology, according to the protocol described by the Earth Microbiome Project, with slight modifications (www.earthmicrobiome.org/emp-standard-protocols). Briefly, frozen extruded feces samples were mechanically disrupted (bead beating), and DNA was extracted with a PowerSoil-htp kit (QIAGEN). From each DNA sample, the 16S rRNA genes from region V3-V4 were PCR-amplified with a composite forward primer and a reverse primer. The reverse primer contained a unique 12-base barcode, designed with the Golay error-correcting scheme, which was used to tag PCR products from respective samples. The composite forward 515F primer sequence was: 5’-*AATGATACGGCGACCACCGAGATCTACACGCT*XXXXXXXXXXXX**TATGGTAATT*GT*** <u>GTGYCAGCMGCCGCGGTAA</u>-3’ where the italicized sequence is the 5’ Illumina adaptor, the 12 X sequence is the golay barcode, the bold sequence is the primer pad, the italicized and bold sequence is the primer linker, and the underlined sequence is the conserved bacterial primer 515F. The reverse primer 806R used was 5’-*CAAGCAGAAGACGGCATACGAGAT***AGTCAGCCAG*CC***<u>GGACTACNVGGGTWTCTAAT</u>-3’ where the italicized sequence is the 3’ reverse complement sequence of Illumina adaptor, the bold sequence is the primer pad, the italicized and bold sequence is the primer linker and the underlined sequence is the conserved bacterial primer 806R. PCR reactions consisted of Hot Master PCR mix (Quantabio, Beverly, MA, USA), 0.2 mM of each primer, 10-100 ng template, and reaction conditions were 3 min at 95°C, followed by 35 cycles of 45 s at 95°C, 60 s at 50°C and 90 s at 72°C on a Biorad thermocycler. PCRs products were quantified using Quant-iT PicoGreen dsDNA assay on a BIOTEK Fluorescence Spectrophotometer and a master DNA pool was generated from the purified products in equimolar ratios. The obtained pool was purified with Ampure magnetic purification beads (Agencourt, Brea, CA, USA), and visualized by gel electrophoresis and then sequenced using an Illumina MiSeq sequencer (paired-end reads, 2×250 bp) at the Genom’IC plateform from Cochin Institut.

### 2.10 16S rRNA gene sequence analysis

16S rRNA sequences were analyzed with QIIME2 – version 2019 360 (Bolyen et al., 2019). Sequences were demultiplexed and quality-filtered with the Dada2 method (Callahan et al., 2016). We used QIIME2 default parameters to detect and correct Illumina amplicon sequence data, and a table of Qiime 2 artifacts was generated. Next, a tree was generated with the align-to-tree-mafft-fasttree command, for analyzing phylogenetic diversity. Then, alpha and beta diversity analyses were computed with the core-metrics-phylogenetic command. We constructed principal coordinates analysis (PCoA) plots to assess the variation between experimental groups (beta diversity). To analyze the taxonomy, we assigned features to operational taxonomic units, according to a 99% threshold of pairwise identity to the Greengenes reference database 13_8. Unprocessed sequencing data are deposited in the European Nucleotide Archive under accession number PRJEB54776, publicly accessible at https://www.ebi.ac.uk/ena/browser/view/PRJEB54776.

### 2.11 Statistical analysis

Data were analyzed with R (http://www.r-project.org). Differential effects were assessed on log2-transformed data by performing analyses of variance (ANOVAs), followed by Student’s t-tests with a pooled variance estimate. P-values from t-tests were adjusted with the Benjamini-Hochberg correction. A p-value <0.01 was considered significant.

We performed hierarchical clustering of gene expression data and lipid quantification data with the R packages, Geneplotter and Marray (https://www.bioconductor.org/). We used Ward’s algorithm, modified by Murtagh and Legendre, as the clustering method. Comparisons were performed with ANOVAs. All data represented on heat maps had p-values <0.05 for one or more comparisons.

Statistical analyses of microbiota data were performed with GraphPad Prism for Windows (GraphPad Prism 7.03). When one-way or two-way ANOVAs found statistically significant differences, they were followed by the appropriate posthoc test (Tukey). Comparisons between two groups were performed with the student’s t-test. P-values <0.05 were considered significant.

## 3. Results

### 3.1 FB1 exposure attenuates the effect of HFD feeding on body weight and fasting glycemia

Eight-week-old C57BL/6J male mice were either fed a low-fat chow diet (10% fat, CHOW) or a HFD (60% fat) *ad libitum* for 15 weeks. At the beginning of the experiment, the four groups of mice were homogeneous in terms of weight. The two groups of mice fed the HFD became overweight within 12 weeks and gained an average of 2g per week per mouse (Fig. S1A). During the same time period, the two groups of mice fed the CHOW diet only gained 4 g body weight (bw) per mouse (Fig. 1A). HFD-fed mice gained significantly more weight, starting from the second week of HFD feeding (Fig. S1B). The difference in body weight continued until the 12th week, when half the mice in each group were exposed to FB1. Thus, during the last three weeks, FB1 (10 mg/kg bw/day) was only added to the drinking water of FB1-exposed groups. From the 12th week to the end of the experiment, FB1 exposure did not affect the weight of CHOW-fed mice, but it induced significant weight loss in HFD-fed mice (around 5 g per mouse; Figure 1A). An evaluation of the food consumed during the last 3 weeks revealed a significant reduction in daily quantity of food intake associated with the HFD-diet in mice exposed to FB1 (but not in the energy intake that significantly increase – Fig. S1C), but FB1 did not significantly influence feeding in CHOW-fed mice (Fig. 1B). In the same period, water consumption increased in mice exposed to FB1 under the CHOW diet, but not in mice under the HFD (Fig. 1C). We checked water consumption to monitor the FB1-exposure level during the experiment and found that exposure to FB1was similar in both dietary groups (HFD = 10.5 ± 0.2 mg/kg bw/day *vs*. CHOW = 10.7 ± 0.6 mg/kg bw/day; Fig. 1D).

**Figure 1.**
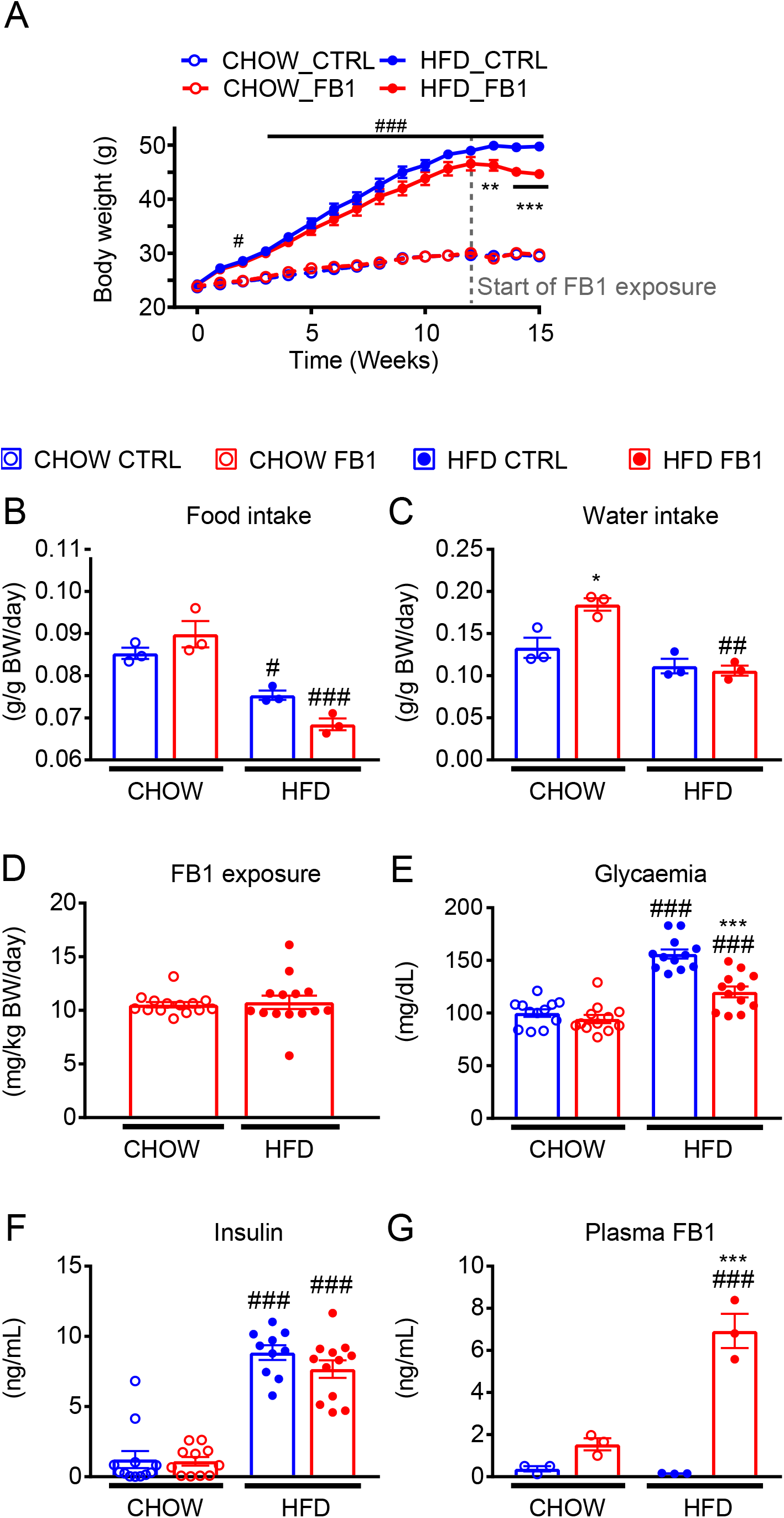
FB1 exposure reverses the effect of HFD on body weight and fasting glucose. C57BL/6J male mice were fed a control diet (CHOW) or a high-fat diet (HFD) for 15 weeks. During the final three weeks, FB1 (10 mg/kg bw/day) was added or not to the drinking water. (A) Mean body weight measured weekly during the study period. (B) Average food intake during the 3 weeks of FB1 exposure. (C) Average water intake during the 3 weeks of FB1 exposure (D) Average FB1 exposure. (E) Fasting glycemia after 2 weeks of FB1 treatment. (F) Insulin levels in the fed state after 3 weeks of FB1 treatment. (G) FB1 level in the plasma. Results are the mean ± SEM (n=12/group and each level correspond to the pooling of 4 mouse samples). # diet effect, * treatment effect. * or # p-value<0.05, ** or ## p-value<0.01, *** or ### p-value<0.001; FB1: fumonisin B1; CTRL: not exposed to FB1

In response to HFD feeding, we observed significant increases in the levels of fasting blood glucose (Fig. 1E) and blood insulin (Fig. 1F). However, FB1-exposed mice under the HFD had significantly lower fasting blood glucose levels than the unexposed HFD-fed mice.

Finally, we evaluated plasma FB1 levels to determine whether the HFD modulated the oral bioavailability of FB1 (Fig. 1G). A comparison between FB1-exposed mice fed CHOW or HFD showed that the HFD increased the FB1 plasma level by 4.5-fold, from 1.54 ± 0.2 ng/ml to 6.92 ± 0.8 ng/ml. Taken together, these results demonstrate that HFD-induced obesity and hyperglycemia blood level were partially reversed by FB1 exposure. This FB1 effect observed in obese mice was correlated with an increase plasma concentration of FB1.

### 3.2 HFD feeding and FB1 exposure influence gut microbiota composition

Next, we investigated whether FB1 effects on obesity and glycemia were related to altered gut homeostasis. We analyzed the effects of both HFD feeding and FB1 exposure on cecal microbial structure through V3-V4 hypervariable regions in 16S rRNA high throughput sequencing. Under a CHOW diet, FB1 exposure did not impacted intestinal microbiota alpha diversity while, as expected, HFD was associated with significant decrease in alpha diversity, as assessed by both the Shannon and Simpson index (Fig. 2A). Importantly, in HFD-fed mice exposed to FB1, alpha diversity was similar to that observed in CHOW-fed mice, suggesting an impact of both HFD and FB1 in regulating intestinal microbiota composition. In order to investigate which phyla were impacted by HFD and/or FB1, we next explored the relative frequencies of taxa at the phylum level (Fig. 2B). HFD feeding significantly decreased the relative frequency of Firmicutes and Actinobacteria and increased the relative frequency of Proteobacteria. In CHOW-fed mice, FB1 did not significantly change the Proteobacteria frequency, but the Actinobacteria and the Firmicutes frequencies were significantly reduced, while the Verrucomicrobia frequency was significantly increased, compared to the frequencies observed in unexposed CHOW-fed mice. In HFD-fed mice, FB1 exposure had little or no significant effects on the relative frequencies of Actinobacteria and Firmicutes. Nevertheless, these results showed that FB1 did not have either synergistic or cumulative effects. For Proteobacteria, the HFD combined with FB1 exposure attenuated the increased relative frequency observed with the HFD alone. However, FB1 induced an increase in the frequency of the Verrucomicrobia phylum in HFD-fed mice.

**Figure 2.**
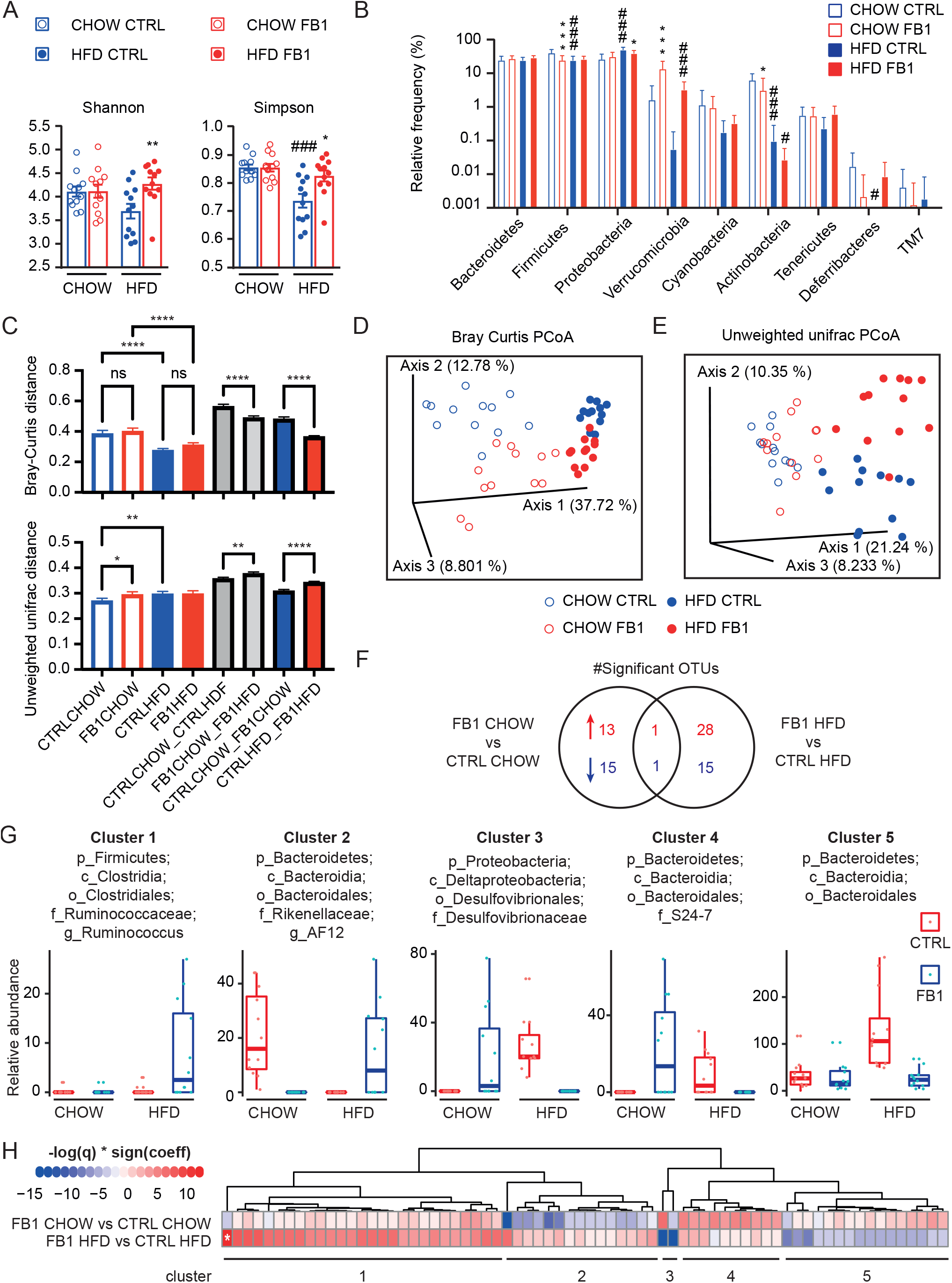
FB1 effects on gut microbiota composition. C57BL/6J male mice were fed a control diet (CHOW) or a high-fat diet (HFD) for 15 weeks. During the final three weeks, FB1 (10 mg/kg bw/day) was added or not to the drinking water. The cecal microbial composition of samples was analyzed by sequencing 16S rRNA genes. (A) Alpha diversity was assessed by calculating the Shannon and Simpson indexes. (B) Relative frequencies of taxa at the phylum level. (C) Beta diversity was assessed with the Bray-Curtis and unweighted unifrac dissimilarity indexes and distances between individuals within and between groups were compared. (D) PCoA plot of beta-diversity using the Bray Curtis index. (E) PCoA plot of beta-diversity using the unweighted unifrac index. (F) General linear models were fitted to find OTUs significantly different between the experimental groups. Venn diagram representing the number of significant OTUs higher (red) or lower (blue) in FB1-vs. CTRL groups. (G) Hierarchical clustering of the OTUs significantly different between FB1 and CTRL mice in either CHOW- or HFD-fed mice. (H) Relative abundance of one representative OTU from each cluster. Data are presented as the mean ± SEM (n=12/group). #diet effect, *treatment effect; * or # p-value<0.05, ** or ## p-value<0.01, *** or ### p-value<0.001; FB1: Fumonisin B1; CTRL: not exposed to FB1; PCoA: principle coordinates analysis

Beta diversity was next evaluated using the Bray-Curtis and unweighted unifrac dissimilarity indexes (Fig. 2C-E). Both PCoA plots showed that HFD feeding was the main factor driving differences in gut microbiota composition, with a clear separation along the 1^st^ PCoA axix. The Bray-Curtis PCoA plot illustrates a significant effect of FB1 in both CHOW- and HFD-fed mice. In the unweighted unifrac PCoA, the FB1-CHOW and the CTRL-CHOW groups were merged, while significant distinct clustering was observed between FB1-HFD and CTRL-HFD groups (Fig 2E), suggesting a stronger impact of FB1 in HFD-fed mice on low abundant ASVs. These findings were confirmed by investigation of the distances separating individual animals within or between groups (Figure 2C). The bray-curtis distance between the FB1- and CTRL-treated animals fed a CHOW diet was significantly lower than the distance between the FB1- and CTRL-treated animals fed a HFD diet, while the opposite pattern was observed using the unweighted unifrac distance (Fig 2D). This indicates that FB1 effects on the gut microbiota seem to depend on the animal diet, with FB1 impacting mostly low abundant bacteria upon HFD feeding.

Finally, we conducted association analysis between microbial ASVs and experimental groups using general linear models (Fig 2F-H). Upon CHOW diet, we found 14 ASVs significantly more abundant, and 16 ASVs significantly less abundant in FB1-treated vs. CTRL mice; while upon HFD diet, 29 ASVs were significantly more abundant, and 16 ASVs significantly less abundant, in FB1-treated vs. CTRL mice. Surprisingly, only 2 ASVs were significantly impacted by FB1 under both dietary regimen (Fig 2F). Adjusted q-value-based hierarchical clustering of these significant OTUs further illustrates this diet-dependent impact of FB1 on gut microbiota, with the ASVs clearly clustering into 5 different clusters (Fig 2G). Among those, ASVs belonging to clusters 1 and 5, illustrate a clear FB1*diet interaction, with FB1 impacting ASVs relative abundance only in HFD-fed mice (Fig 2H).

Taken together, these results provide evidence that the ecological balance of gut microbiota was significantly modified by both the HFD and FB1 exposure. Moreover, we observed an interaction between HFD and FB1 on the intestinal microbiota composition.

### 3.3 FB1 reverses HFD-induced hepatic steatosis, but promotes liver inflammation

Next, we performed histological analyses of the liver to assess the effects of HFD feeding and FB1 exposure on liver physiology and homeostasis (Fig. 3A). Histological H&E staining showed that HFD feeding induced steatosis. In CHOW-fed mice, FB1 exposure did not induce any detectable morphological differences from unexposed samples. However, in the HFD group, FB1 exposure induced a marked reduction in steatosis compared to the unexposed group. These results were associated with a significant decrease in liver weight (Fig. 3B), steatosis scores (Fig. 3C), hepatic triglycerides (Fig. 3D) and in some mRNA relative genes expression corresponding to lipogenesis (Fig. S2A). Additionally, both hepatic free-cholesterol and esterified cholesterol were increased in the HFD group compared to the CHOW group, but FB1 exposure did not significantly affect these HFD effects (Fig. 3E and 3F).

**Figure 3.**
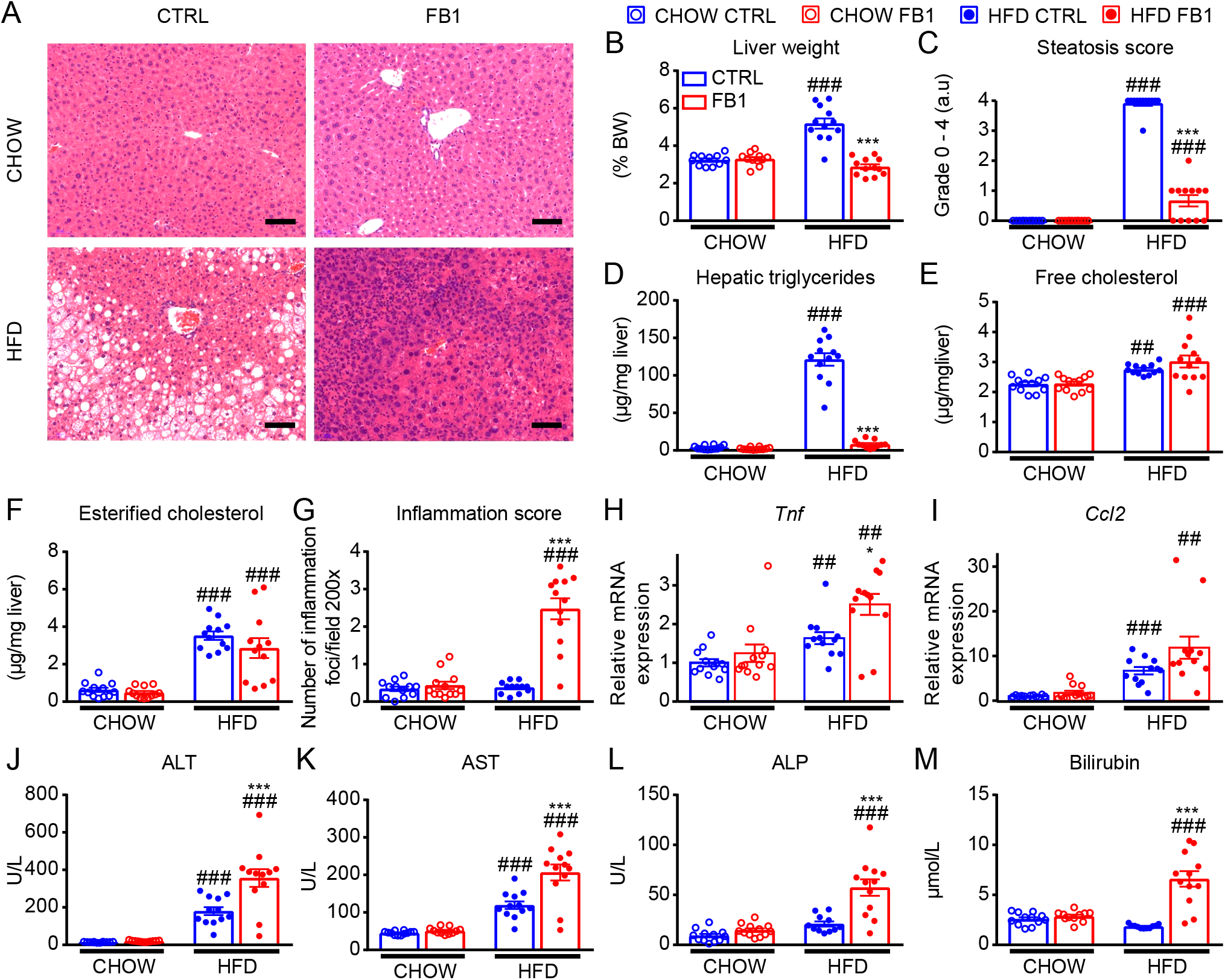
FB1 reverses HFD-induced hepatic steatosis, but promotes liver inflammation. C57BL/6J male mice were fed a control diet (CHOW) or a high-fat diet (HFD) for 15 weeks. During the final three weeks, FB1 (10 mg/kg bw/day) was added or not to the drinking water. (A) Representative histological liver sections from mice in each group stained with hematoxylin and eosin; magnification ×100. Scale bars: 50 µm. (B) Average liver weight, expressed as a percentage of body weight. (C) Liver steatosis scores, estimated on liver sections. Scoring: the severity of parenchymal steatosis depended on the percentage of liver cells that contained fat: Grade 0: no hepatocytes with steatosis in any section; grade 1: 1−25% of hepatocytes with steatosis; grade 2: 26−50% of hepatocytes with steatosis; grade 3: 51−75% of hepatocytes with steatosis, and grade 4: 76−100% of hepatocytes with steatosis (n=12/group). (D-F) Lipids were extracted from livers and quantified with gas-liquid chromatography: (D) triglycerides, (E) free cholesterol, and (F) esterified cholesterol. (G) Inflammatory scores: liver sections were analyzed in 10 microscopic fields (200× magnification) to determine the mean number of inflammatory foci per field (n=12 per group). (H-I) mRNA expression levels of genes that encode cytokines involved in inflammation: (H) *Tnfα* and (I) *Ccl2.* (J-M) End of experiment plasma levels of (J) aspartate aminotransferase (AST), (K) alanine aminotransferase (ALT), (L) alkaline phosphatase (ALP), and (M) bilirubin. Results are presented as the mean ± SEM. #diet effect, *treatment effect; * or # p-value< 0.05, ** or ## p-value<0.01, *** or ### p-value<0.001; FB1: Fumonisin B1; CTRL: not exposed to FB1

Furthermore, H&E staining revealed that liver sections from mice fed the HFD and exposed to FB1 had significantly more inflammatory foci than any of the other mouse groups (Fig. 3A). Liver inflammation was confirmed by the inflammatory score (Fig. 3G), and by the significant increases in the relative hepatic expression of *Tnf* and *Ccl2* mRNA (Fig. 3H,I). Although both of these genes were significantly upregulated in response to the HFD, only the relative expression *Tnf* mRNA was significantly increased with FB1 exposure, compared to HFD feeding alone (Fig. S2B,C).

Liver damage was confirmed by analyzing plasma levels of ALT (Fig. 3J) and AST (Fig. 3K). Both these enzymes were elevated in HFD-fed mice compared to CHOW-fed mice. In HFD-fed mice, FB1 exposure caused further elevations of ALT and AST. In addition, the plasma ALP and total bilirubin levels were significantly increased when HFD-fed mice were exposed to FB1 (Fig. 3L,M).

Taken together, these data suggest that the HFD combined with FB1 reversed HFD-induced hepatic steatosis, but promoted liver inflammation and hepatocytolysis. The HFD combined with FB1 perturbed the bile and bilirubin.

### 3.4 Effect of FB1 on hepatic sphingolipid homeostasis

With FB1 being a known ceramide synthase inhibitor, we next investigated FB1-induced alterations in hepatic sphingolipid metabolism in both CHOW-fed and HFD-fed mice. We measured several sphingolipid species in the liver, including sphinganine (Sa), sphingosine (So), sphingosine-1-phosphate (S1P), ceramides, dihydroceramides, and sphingomyelins (Fig. 4).

**Figure 4.**
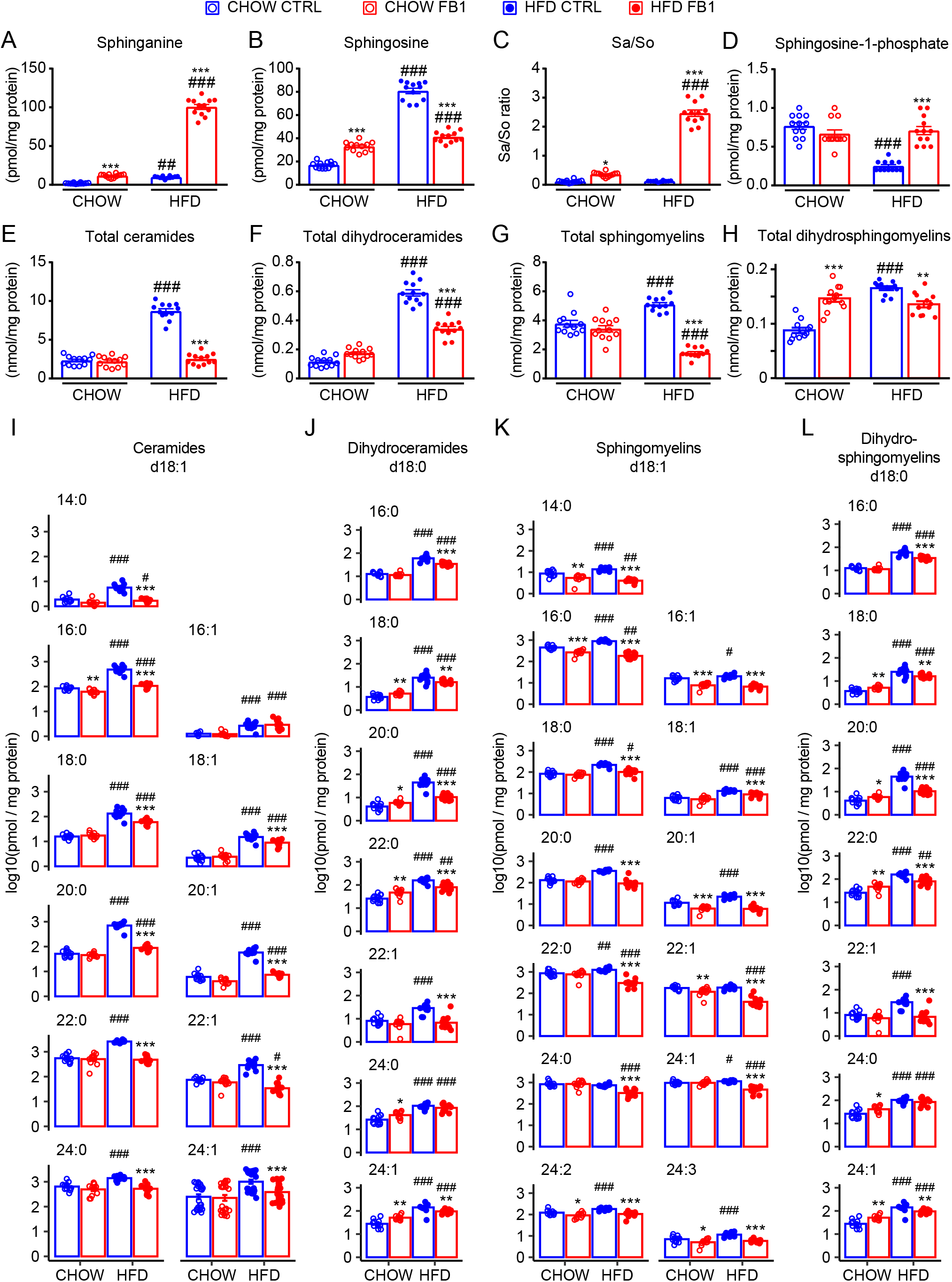
FB1 effects on sphingolipid homeostasis. C57BL/6J male mice were fed a control diet (CHOW) or a high-fat diet (HFD) for 15 weeks. During the final three weeks, FB1 (10 mg/kg bw/day) was added or not to the drinking water. We analyzed liver samples for the levels of (A) sphinganine, (B) sphingosine, (C) the sphinganine/sphingosine ratio (Sa/So), (D) sphingosine-1-phosphate, (E) total ceramides, (F) dihydroceramides, (G) sphingomyelins, and (H) dihydrosphingomyelins. (I-L) To evaluate the abundances of sphingolipids as a function of the length of the fatty acid residue, we performed separate measurements of (I) ceramide, (J) dihydroceramide, (K) sphingomyelin, and (L) dihydrosphingomyelin species. Results are presented as the mean ± SEM. #diet effect, *treatment effect. * or # p-value< 0.05, ** or ## p-value<0.01, *** or ### p-value<0.001; FB1: Fumonisin B1; CTRL: not exposed to FB1

As expected, under the CHOW diet, FB1 exposure induced significant increases in the hepatic levels of sphingoid bases and of the Sa/So ratio (3-fold increase). These sphingoids are well-known biomarkers for FB1 effects (Fig. 4A-C). Moreover, the total hepatic levels of dihydrosphingomyelins also increased significantly with FB1 exposure under the CHOW diet (Fig. 4H). Surprisingly, under the CHOW diet, the level of FB1 exposure applied did not significantly affect the hepatic levels of S1P, total ceramides, total dihydroceramides, or total sphingomyelins (Fig. 4D-G). A closer look at the specific ceramide and sphingomyelin species (Fig. 4I, 4K) showed that the abundances of some were significantly reduced, including ceramide(d18:1/16:0), sphingomyelin(d18:1/14:0), sphingomyelin(d18:1/16:0), sphingomyelin(d18:1/16:1), sphingomyelin(d18:1/20:1), sphingomyelin(d18:1/22:1), sphingomyelin(d18:1/24:2), and sphingomyelin(d18:1/24:3). Moreover, under the CHOW diet, FB1 exposure induced significantly higher levels of specific dihydroceramides (Fig. 4J) and long carbon-chain dihydrosphingomyelins (Fig. 4L).

Under HFD feeding, the basal hepatic levels of ceramides, dihydroceramides, sphingomyelins, and dihydrosphingomyelins significantly increased (Fig. 4E-H). Similarly, the levels of sphinganine and sphingosine increased, but the Sa/So ratio remained unchanged (Fig. 4A-C). In contrast, the level of S1P significantly decreased with the HFD (Fig. 4D). Analyzing the specific ceramides, dihydroceramides, sphingomyelins, and dihydrosphingomyelins species, we found that HFD feeding caused significant elevations in nearly all species (Fig. 4I-L).

When the HFD was combined with FB1 exposure, stronger effects were observed on sphingolipid metabolism. This combined treatment induced a significant increase in the hepatic sphinganine levels (Fig. 4A) and a reduction in the hepatic sphingosine levels, to the level observed in unexposed HFD-fed mice, but not to the level observed in CHOW-fed unexposed mice (Fig. 4B). These changes in sphingoid base levels resulted in a marked increase in the Sa/So ratio (20-fold increase), which is characteristic of severe FB1 contamination (Fig. 4C). Moreover, when the HFD was combined with FB1 exposure, the reduced sphingosine level was associated with a significant increase in the S1P level, to the level observed in unexposed HFD-fed mice (Fig. 4D). In addition, the HFD combined with FB1 exposure caused significant reductions in the hepatic levels of ceramide, dihydroceramides, sphingomyelins, and dihydrosphingomyelins, compared to unexposed HFD-fed mice (Fig. 4E-L). Nevertheless, the total sphingomyelin level was reduced to a significantly lower level than that observed in unexposed CHOW-fed mice, the total ceramide level was reduced to the same level as that observed in unexposed CHOW-fed mice. Finally, in HFD-fed mice exposed to FB1, the dihydroceramide and dihydrosphingomyelin levels remained significantly higher than the levels observed in unexposed CHOW-fed mice.

Taken together, these results suggest that HFD-induced liver steatosis enhance FB1 effect on sphingolipid metabolism inhibiting more efficiently ceramide synthetase. Surprisingly, under HFD-induced obesity FB1 seems to enhance sphingosine-kinase activity and prevent glycosphingolipid recycling (Fig. S3).

### 3.5 Effect of FB1 on the hepatic metabolome

These severe metabolic effects on sphingolipids and the previous reports (Régnier *et al*., 2017 ; Régnier *et al*., 2019b) that indicated that an HFD combined with FB1 exposure had an impact on lipid metabolism encouraged us to explore the global metabolomic profile of the liver with an untargeted approach.

To investigate the effect of FB1 on hepatic metabolism, we performed ^1^H-NMR-based metabolic profiling on liver tissues. We generated O-PLS-DA plots derived from ^1^H-NMR spectra of aqueous hepatic extracts and compared the effects of FB1 exposure on the liver metabolic profile under either CHOW or HFD feeding. No significant effects of FB1 exposure on the profiles of CHOW-fed mice were observed (Fig. 5A). However, FB1 exposure left a clear, significant metabolic fingerprint in HFD-fed mice (Fig. 5B). The coefficient plot derived from the O-PLS-DA model for HFD-fed mice highlighted differences in the levels of particular metabolites associated with FB1-exposure (Fig. 5C). For example, FB1 exposure specifically impacted the ^1^H-NMR chemical shift signals of bile acids, glutamate, succinate, aspartate, dimethylamine, taurine, choline, glycerophosphocholine (GPC), fumarate, tyrosine, and uridine.

**Figure 5.**
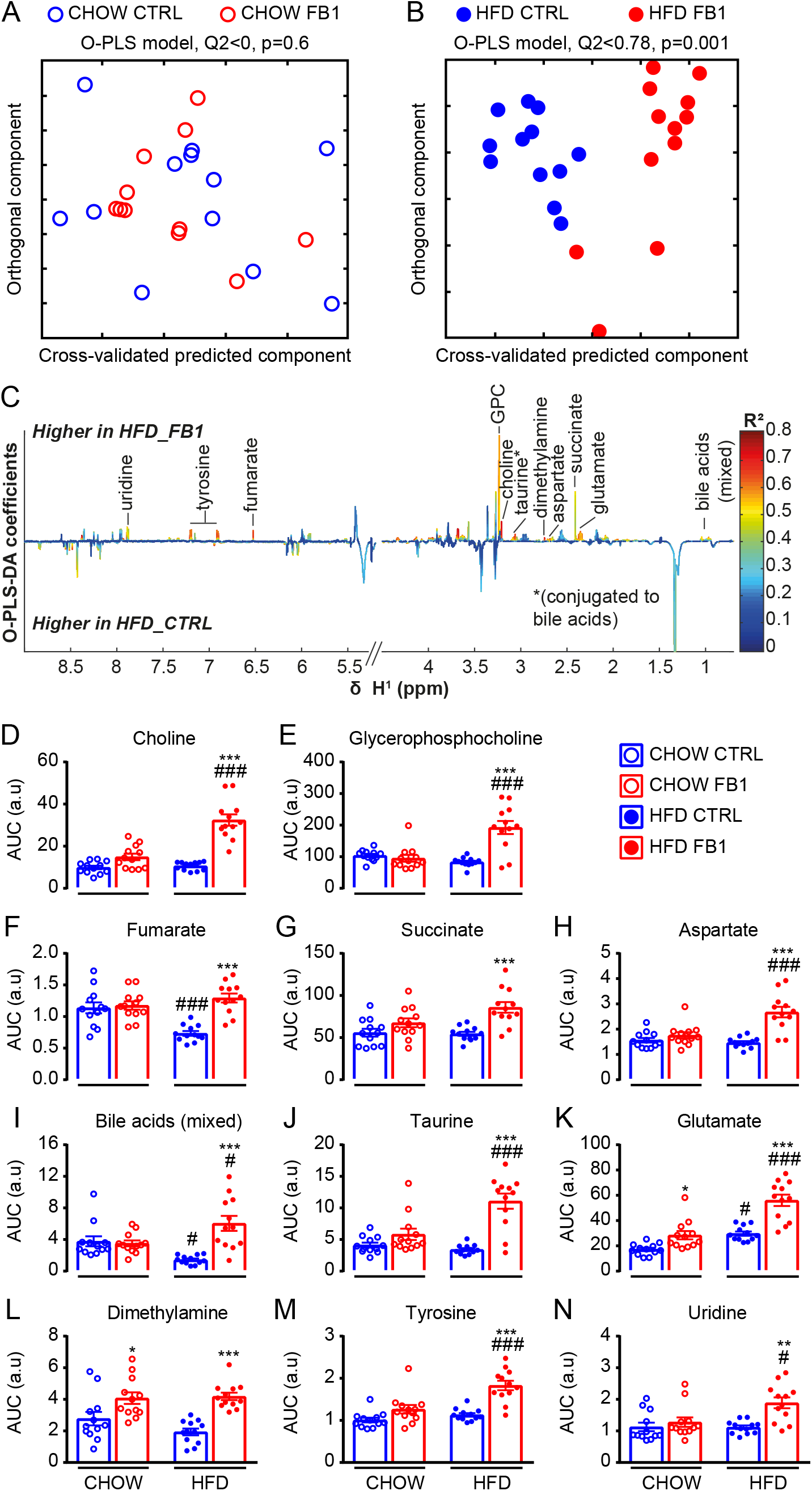
FB1 effects on the metabolomic profile of the liver. C57BL/6J male mice were fed a control diet (CHOW) or a high-fat diet (HFD) for 15 weeks. During the final three weeks, FB1 (10 mg/kg bw/day) was added or not to the drinking water. (A-B) O-PLS-DA score plots derived from the ^1^H-NMR metabolomic profiles of liver aqueous extracts from CHOW (A) or HFD (B)-fed mice. Each dot represents an animal. (C) Coefficient plots related to the O-PLS-DA models discriminating between HFD alone (HFD_CTRL) and HFD combined with FB1 exposure (HFD_FB1). Metabolites are color-coded according to their correlation coefficient. The direction of the metabolite peak indicates the group with which it was positively associated, as labeled on the diagram. (D-N) Areas under the curves for several discriminant metabolites selected using the previous O-PLS-DA model. Additional 2-way ANOVAs confirmed significant differences in metabolite levels (n=12/group). Results are presented as the mean ± SEM. #diet effect, *treatment effect. * or # p-value< 0.05, ** or ## p-value<0.01, *** or ### p-value<0.001; FB1: Fumonisin B1; CTRL: not exposed to FB1; O-PLS-DA: orthogonal projection on latent structure-discriminant analysis; GPC: glycerophosphocholine.

The areas under the curves of the ^1^H-NMR spectra were integrated for metabolites that were significantly correlated with the predictive component (R^2^>0.5). Univariate statistics (1-way ANOVA + Sidak’s post-tests) confirmed significant increases in the levels of metabolites involved in choline metabolism (choline and glycerophosphocholine); the tricarboxylic acid cycle (fumarate, succinate, aspartate, and glutamate); biliary acid metabolism (mixed bile acids and taurine); intestinal microbiota dysbiosis (dimethylamine and tyrosine), and uridine metabolism (Fig. 5D-N). These metabolic profile analyses confirmed that obesity induced by HFD feeding significantly influenced the effect of FB1 exposure on liver metabolism *in vivo*.

### 3.6 Effect of FB1 exposure on liver gene expression

We next performed an unbiased microarray analysis of liver gene expression to identify biological processes that were sensitive to FB1 exposure under both CHOW and HFD feeding. A principal component analysis (PCA) of the transcriptome showed a clear separation between CHOW-fed and HFD-fed groups (Fig. 6A). The separation observed along the second axis accounted for 13.6% of the variance. Upon CHOW-fed, the unexposed and FB1 exposed groups overlapped. In contrast, the unexposed and FB1 exposed HFD-fed groups were clearly separated. The separation along the first axis accounted for 56.3% of the variance.

**Figure 6.**
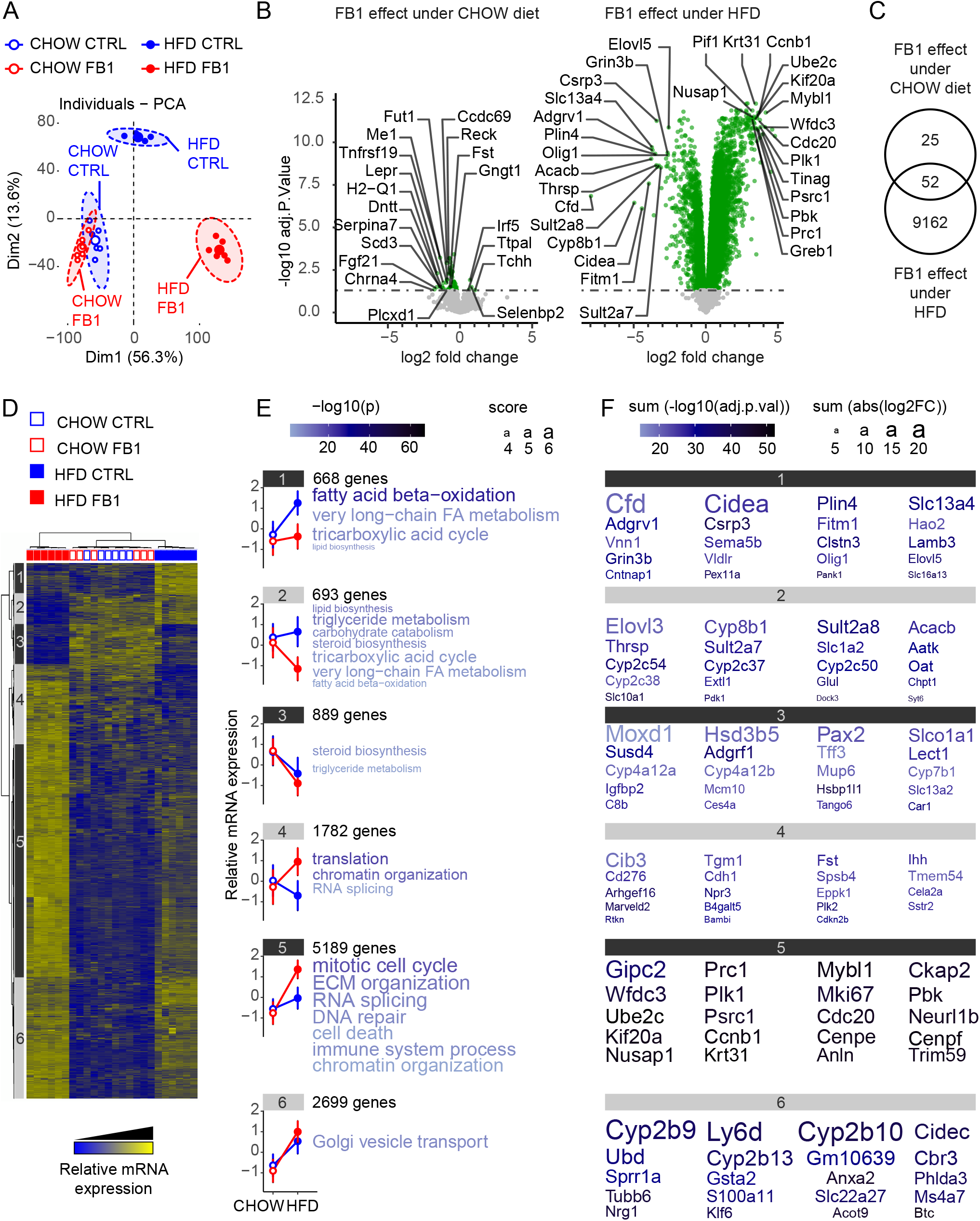
FB1 effects on liver gene expression. C57BL/6J male mice were fed a control diet (CHOW) or a high-fat diet (HFD) for 15 weeks. During the final three weeks, FB1 (10 mg/kg bw/day) was added or not to the drinking water. Gene expression profiles were analyzed in liver samples with Agilent microarrays (n=6/group). (A) Principal component analysis (PCA) score plots of whole-liver transcriptome datasets (n=6/group). Each dot represents an observation (animal), projected onto first (horizontal axis) and second (vertical axis) PCA variables. (B) Volcano plot shows FB1 effects on gene expression under a CHOW diet (left panel) or an HFD (right panel). Each gene expression level is shown in terms of the −log10 p-value, for comparisons between the FB1 exposed group and the unexposed (CTRL) group for each diet. The −log10 p-values are plotted as a function of the associated log2-fold change, or formally, log2(FB1)-log2(CTRL). The green points have p-values <0.05. Gene names are highlighted for the most highly regulated genes, according to a score based on the adjusted p-value × logFC. (C) Venn diagram represents the number of genes significantly regulated by FB1 exposure for each diet. (D) Heatmap represents data from microarray experiments. The significantly differentially expressed genes (adjusted p-values <0.05) were selected, and they corresponded to 11,920 probes. The color gradient indicates the scaled values of gene expression. Hierarchical clustering identified six gene clusters (indicated on the left). (E) Mean expression profiles for the six gene clusters. Graphs represent the means of the scaled gene expression values. Error bars are standard deviations. The most significantly enriched biological processes identified with the Metascape gene ontology algorithm are shown at the right of each profile. Briefly, hypergeometric tests were performed for each category in each cluster. The size of the font is related to a score based on the log base 2 number of genes enriched, and the color gradients of the characters represent the −log base 10 value of the probability of the test for P[X > x]. (F) Representation of the top 20 genes in each cluster that showed the largest differences in expression. The color of each character string is related to the sum of the −log10(adjusted p-value), and the size of each character string is related to the sum of the absolute log2FC values for all the comparisons made for each gene.

Volcano plots of the FB1 effect upon CHOW-fed or HFD-fed mice confirmed the stronger genomic response to FB1 exposure with HFD feeding (Fig. 6B). Indeed, only 77 genes showed significantly modulated expression with FB1 exposure under the CHOW diet. In contrast, with the HFD, 9,214 hepatic genes were differentially expressed in response to FB1 exposure (Fig. 6C).

We then performed hierarchical clustering to analyze the differentially expressed genes (those with adjusted p-values<0.05), which corresponded to 11,920 probes (Fig. 6D). Along the horizontal axis, the blind clustering of profiles did not discriminate between FB1 exposed and unexposed mice under the CHOW diet. Conversely, HFD feeding induced marked clustering that discriminated clearly between unexposed mice and FB1 exposed mice. An analysis of the gene clustering revealed 6 major genetic groups along the vertical axis of the heatmap (Fig. 6D). Of these, four clusters were related to genes with similar expression levels in FB1-exposed mice under the CHOW diet but differentially expressed genes in FB1-exposed mice under HFD diet.

Expression of genes from clusters 1 and 2 was reduced upon FB1 exposure in HFD-fed mice. These genes were related to energy metabolism. In the first cluster, 668 genes showed an important increase in mRNA expression under the HFD compared to the CHOW diet. However, when the HFD group was exposed to FB1, mRNA expression was similar to the levels observed under the CHOW diet, with or without exposure to FB1. Moreover, the gene ontology enrichment analysis of this set of genes (Fig. 6E) revealed that the biological processes most significantly associated with this cluster were: fatty acid beta-oxidation, very long-chain fatty acid metabolism, and the tricarboxylic acid cycle. Furthermore, characterization of the most significantly affected genes in cluster 1 (Fig. 6E) showed that, under HFD feeding, FB1 exposure essentially limited increases in the expression of genes involved in triglyceride storage, such as *Cidea, Fitm1, Plin4, Vldlr*, and *Elovl5*. In contrast, the 693 genes in cluster 2 showed an important reduction in mRNA expression under HFD feeding with FB1 exposure, compared to the CHOW-fed, unexposed group. Moreover, under HFD-feeding alone, mRNA expression was similar to the levels observed under the CHOW diet, with or without FB1 exposure. Similar to cluster 1, the gene ontology enrichment analysis of this set of genes (Fig. 6D) revealed that the biological processes most significantly associated with cluster 2 were: triglyceride metabolism, the tricarboxylic acid cycle, very long-chain fatty acid metabolism, carbohydrate catabolism, and steroid biosynthesis. Furthermore, characterization of the most significantly affected genes in cluster 2 (Fig. 6E) showed that, under HFD feeding, FB1 exposure reduced expression of genes involved in fatty acid metabolism, such as: *Elovl3* (involved in very long-chain fatty acid elongation from C18:0 to provide precursors for sphingolipid synthesis); *Acacb* and *Pdk1* (involved in fatty acid uptake and oxidation in mitochondria); and *Thrsp* (involved in lipid storage).

Clusters 4 and 5 included genes involved in cell cycle metabolism and organization. Indeed, the expression levels of the 1,782 genes in cluster 4 were slightly decreased under the HFD, compared to the CHOW diet. However, a moderate increase in mRNA expression was observed with the HFD and FB1 exposure, compared to the CHOW diet with FB1 exposure. Moreover, the gene ontology enrichment analysis (Fig. 6D) revealed that the biological processes most significantly associated with cluster 4 were translation, chromatin organization, and RNA splicing. Furthermore, characterization of the most significantly affected genes in cluster 4 (Fig. 6E) showed that, under HFD feeding, FB1 exposure reversed and slightly increased the expression of genes involved in cell proliferation (*Tgm1, Eppk1*) and cell junction organization (*Marveld2, Cdh1*). In cluster 5, the expression of 5,189 genes significantly increased with the HFD and even more upon FB1 exposure, compared to the CHOW groups, without or with FB1 exposure. This effect indicated synergy between FB1 exposure and the HFD. The gene ontology enrichment analysis (Fig. 6D) revealed that the biological processes most significantly associated with cluster 5 were the mitotic cell cycle, extracellular matrix organization, RNA splicing, DNA repair, immune system processes, chromatin organization, and cell death. Furthermore, characterization of the most significantly affected genes in cluster 5 (Fig. 6E) showed that, under HFD feeding, FB1 exposure significantly amplified the expression of genes involved in cell cycle regulation (*Plk1, Prc1, Ube2c, Cdc20, Ccnb1, Cenpf, Cenpe*) and cytoskeleton organization (*Ckap2, Kif20a, Nusap1, Anln*).

Finally, clusters 3 and 6 exhibited significant modulations with diet, independent of FB1 exposure. Indeed, in cluster 3, the expression levels of 889 genes associated with steroid biosynthesis or triglyceride metabolism decreased significantly under HFD feeding. In contrast, in cluster 6, the expression levels of 2,699 genes associated with Golgi vesicle transport increased under the HFD.

## 4. Discussion

Environmental exposure to natural toxicants or chemical residues, alone or in mixtures, are frequently associated with the risk of chronic metabolic diseases (Grün et al., 2006). Moreover, the increasing prevalence of obesity (Estes et al., 2018), increases the risks of various diseases, including liver injuries. Several studies previously reported that toxicants, like triclosan (Yueh et al., 2020), 2,3,7,8-tetrachlorodibenzo-p-dioxin (Duval et al., 2017), chlorpyrifos (Wang et al., 2021), or methyl tert-butyl ether (Tang et al., 2019) contributed to the progression of obesity-associated liver steatosis. Those findings led us to hypothesize that environmental toxins may differentially impact liver homeostasis, depending on the presence of obesity. Among the natural food contaminants, some of the most prevalent and harmful mycotoxins are known to induce liver toxicity, such as aflatoxin B1 (Fan et al., 2021; Hua et al., 2020; Plaz Torres et al., 2020), T-2 toxin (Janik et al., 2021), deoxynivalenol (Hasuda et al. 2022), ochratoxin A (Tao et al., 2018), zearalenone (Wang et al., 2019), and FB1 (Wangia-Dixon et al., 2021).

It is well-established that FB1 affects the gut-liver axis and liver metabolism (Terciolo et al., 2019; Régnier et al., 2017). Therefore, we tested the toxic effects of FB1 exposure in mice with diet-induced obesity. First, as expected, we showed that HFD feeding induced obesity, glucose intolerance, and hepatic steatosis (Régnier et al., 2020; Tamura et al., 2005). Second, we confirmed the known effect of FB1 exposure on sphingolipid homeostasis, which resulted in an increase in the Sa/So ratio (Régnier et al., 2019; Loiseau et al., 2015). Then, we observed that HFD-induced obesity followed by 4 weeks of co-exposure to an HFD and FB1 resulted in gut dysbiosis, increased plasma FB1 levels, and reductions in body weight, liver weight, fasting blood glucose, and triglyceride levels. However, several plasma markers of liver injury (ALT, AST, ALP, and bilirubin) were significantly increased, which indicated severe hepatitis. Finally, unbiased analyses of the liver metabolome and transcriptome produced results consistent with the notion that FB1 exposure had a potent effect on liver metabolism, which is additive to the effects of diet-induced obesity.

Several lines of evidence have suggested that environmental toxicants may influence obesity and NAFLD (Rajak et al., 2021). However, most preclinical studies supporting this hypothesis were co-exposure studies. In contrast, the present study took an original approach by exposing mice to FB1 after they became obese and hyperglycemic on the HFD. We monitored body weight and water intake to ensure that chow-fed and HFD-fed mice were exposed to a similar dose of FB1 relative to body weight. Thus, with similar FB1 dosing, normal and obese mice showed different systemic and hepatic responses to FB1. However, the plasma FB1 levels were different in CHOW and HFD groups. This result might be due to either increased FB1 absorption or reduced FB1 clearance in the HFD-fed mice.

This study had some limitations. First, our study design did not allow us to determine the mechanism by which HFD exposure induces the increase in circulating plasma level of FB1. The HFD might have changed the gut physiology, altered the microbiota composition and/or activity (Rohr et al., 2020; Mouries et al., 2019), or suppressed FB1 detoxication; indeed, both obesity and hepatic steatosis are known to hamper detoxification processes in the gut and liver (Cobbina *et al*., 2017; Sharpton *et al*., 2019). Another limitation of the study was that we administered a high dose of FB1, which was hundred times above the BMDL_10_ of 0.1 mg/kg bw per day calculated by the CONTAM Panel from EFSA and derived for induction of megalocytic hepatocytes in mice (Bondy *et al*., 2012; Knutsen *et al*., 2018a). Thus, one might question the potential relevance of the findings to animal and human populations (Terciolo et al., 2019). However, rodents are known to be particularly resistant to FB1 toxicity; indeed, very few biological markers have been modulated in rodents under a regular CHOW diet. Nevertheless, the additive effects of HFD feeding and FB1 exposure observed in this study provided further evidence that obesity could weaken the host’s ability to cope with food toxins, and revealed novel insights on the hepatic toxicity of FB1.

In obesity, the liver is exposed to increase in both endotoxin levels and metabolic stress. Both these factors promote NAFLD, which ranges in severity, from steatosis to steatohepatitis, cirrhosis, and cancer (Ferro et al., 2020; Todoric et al., 2020; Loo et al., 2017; Kübeck et al., 2016). Based on our histological analyses and our targeted assays on liver composition and function, we concluded that FB1 reduced the steatosis and neutral lipid deposition induced by HFD feeding. These effects were associated with reductions in body weight and hyperglycemia, which suggested that FB1 could reduce obesity and diabetes, which in turn, might have contributed to reducing hepatic lipid accumulation (Meikle et al., 2017; Holland et al., 2008). Our monitoring of food intake showed that mice did not significantly reduce food intake during FB1 exposure. This result suggested that FB1 affected calorie absorption and/or expenditure. However, this hypothesis warrants future study, because it is beyond the direct effects of FB1 on hepatic homeostasis. Although FB1 exposure reduced steatosis in HFD-fed mice, it also significantly induced liver inflammation, damage, and dysfunction. Indeed, FB1-induced hepatitis was much more severe in HFD-fed mice than in CHOW-fed mice, and it was associated with a massive shift in liver metabolism and gene expression.

It remains unclear whether all of these HFD-exacerbated signs of FB1 toxicity were related to FB1 inhibition of sphingolipid synthesis or whether it involved multiple organ cross-talk. Sphingolipids, such as ceramides, are bioactive lipids that drive the progression of steatosis (Hannun et al., 2018; Choi et al., 2015; Xia et al., 2015). Indeed, several studies have identified correlations between ceramides and different measures of NAFLD in humans. Additionally, various preclinical studies in rodents have demonstrated that ceramides are necessary for NAFLD development (Poss et al., 2020; Chaurasia et al., 2016; Régnier et al., 2019). Therefore, the effects of FB1 that we observed on steatosis were consistent with an inhibition of the steatogenic role of ceramides (Chaurasia et al., 2019; Holland et al., 2008). Furthermore, the effects of FB1 on liver damage and inflammation were consistent with an inhibition of the well-known pro-inflammatory and pro-apoptotic effects of sphingolipid species respectively such as S1P and sphingoïd bases (Molino et al., 2017; Riley et al., 2001). Therefore, the pro-inflammatory effects of FB1 observed in HFD-fed mice might have occurred as an indirect consequence of altered ceramide homeostasis (Chen et al., 2021). Alternatively, high FB1 exposure may exert toxic effects in hepatocytes that are independent of ceramide metabolism, but reflect a mechanism yet to be determined.

## 5. Conclusion

To our knowledge, the present study was the first to assess the effects of diet-induced obesity on FB1 toxicity. This work established that, in an obese context, FB1 exposure exhibited enhanced gut dysbiosis, systemic and hepatotoxic effects. Although FB1 exposure in diet-induced obese mice led to significant reductions in body weight, glycemia, and hepatic lipid content, it also led to liver inflammation and increases in various markers of hepatotoxicity. Therefore, our findings suggested that diet-induced obesity might increase the sensitivity to environmental toxins.

## Declaration of Competing Interest

The authors declare no competing financial interests or personal relationships that could have influenced the study.

## Acknowledgements

L.D. PhD was supported by the INRAE Animal Health department. This work was also supported by grants from the French National Research Agency (ANR) Fumolip (ANR-16-CE21-0003) and the Hepatomics FEDER program of Région Occitanie. We thank Prof Wentzel C. Gelderblom for generously providing the FB1 and for his interest and support in our project. B.C. laboratory is supported by a Starting Grant from the European Research Council (ERC) under the European Union’s Horizon 2020 research and innovation program (grant agreement No. ERC-2018-StG-804135), a Chaire d’Excellence from IdEx Université de Paris - ANR-18-IDEX-0001, an Innovator Award from the Kenneth Rainin Foundation, an ANR grant EMULBIONT ANR-21-CE15-0042-01 and the national program “Microbiote” from INSERM. We thank Anexplo Genotoul MetaToul for their excellent work in plasma biochemistry. Neutral Lipids MS and NMR experiments were performed with instruments in the Metatoul-AXIOM platform. Sphingolipid MS analysis were performed with instruments in the RUBAM platform. The FB1 plasma levels were determined using an UPLC-MS/MS instrument part of the Ghent University MSsmall expertise centre for advanced mass spectrometry analysis of small organic molecules. We thank Elodie Rousseau-Bacquié and all members of the EZOP staff for their assistance in the animal facility. We are very grateful to Talal al Saati for histology analyses and review, and we thank all members of the US006/CREFRE staff at the histology facility and the Genom’IC platforms (INSERM U1016, Paris, France) for their expertise.

## Notes

https://www.ebi.ac.uk/ena/browser/view/PRJEB54776

https://www.ncbi.nlm.nih.gov/geo/query/acc.cgi?acc=GSE208735

### Competing Interest Statement

The authors have declared no competing interest.

### Summary of Updates

author affiliations updated;

