## Supplemental for "Obesity promotes Fumonisin B1 toxicity and induces hepatitis"

**Supplementary Table 1. Oligonucleotides sequences for real-time PCR.**

| GeneName | Forward primer (5'-3') | Reverse primer (5'-3') |
|---|---|---|
| *Acca* | TTACAGGATGGTTTGGCCTTTC | CAAATTCTGCTGGAGAAGCCAC |
| *Acly* | AAAGCTTGGCCTCGTCGG | GGGACGAAGGGTTCAATGAGA |
| *Casp1* | GCTTCAATCAGCTCCATCAGC | CAGTCCTGGAAATGTGCCAT |
| *Ccl2* | GGTGTCCCAAAGAAGCTGTAGTTTT | AGTTGTAGGTTCTGATCTCATTTGGTT |
| *Col3a1* | TGGCACAGCAGTCCAACGTA | CATAGGACTGACCAAGGTGGCT |
| *Elovl6* | TCTGATGAACAAGCGAGCCA | TGGTCATCAGAATGTACAGCATGT |
| *Fasn* | AGTCAGCTATGAAGCAATTGTGGA | CACCCAGACGCCAGTGTTC |
| *Gck* | TCGCAGGTGGAGAGCGA | TCGCAGTCGGCGACAGA |
| *Krt18* | CAAGTACTGGTCTCAGCAGATTGA | CTTGGTGGTGACAACTGTGGTA |
| *Lpk* | TCGACTCAGAGCCTGTGGC | AGTCGTGCAATGTTCATCCCT |
| *Myd88* | CTGGCCTTGTTAGACCGTGAG | TGGTTCTGCTGCTTACCTAAGTATT |
| *Nlrp3* | AGCTAAGAAGGACCAGCCAGA | GGCATACCATAGAGGAATGTGATGTA |
| *Panx1* | TGGCCTTCGCTCAGGAGAT | CAGCAGCCCAGCAGTATGAAT |
| *Pycard* | GCTGCAAACGACTAAAGAAGAGTC | GTGTCCTGTTCTGGCTGTACTC |
| *Scd1* | CAGTGCCGCGCATCTCTAT | CAGCGGTACTCACTGGCAGA |
| *Srebp1-c* | GGAGCCATGGATTGCACATT | GCTTCCAGAGAGGAGGCCAG |
| *Tnf* | TCCCCAAAGGGATGAGAAGTTC | GCGCTGGCTCAGCCACT |
| *Tlr4* | CCTGACACCAGGAAGCTTGAAT | TCATCAGGGACTTTGCTGAGTTT |

### SUPPLEMENTARY FIGURE LEGENDS

**Figure S1. FB1 exposure reverses the effect of HFD on body weight and fasting glucose.**

C57BL/6J male mice were fed a control diet (CHOW) or a high-fat diet (HFD) for 15 weeks. During the final three weeks, FB1 (10 mg/kg bw/day) was added or not to the drinking water. (A) Mean of the weekly body weight gain along the 12 first weeks before FB1 exposure. Results are the mean ± SEM (n=12 weeks/group and each level correspond to the pooling of data from 24 mice per group). (B) Mean of the weekly body weight gain along the 3 weeks of FB1 exposure. Results are the mean ± SEM (n=3 weeks/group and each level correspond to the pooling of data from 12 mice per group). (C) Average of the daily energy food intake along the 15 weeks of experiments. Results are the mean ± SEM (n=number of weeks/group and each level correspond to the pooling of data from 12 mice per group). # diet effect, * treatment effect. * or # p-value<0.05, **** p-value<0.0001; FB1: fumonisin B1; CTRL: not exposed to FB1.

**Figure S2. Effect of FB1 on liver gene expression.**

C57BL/6J male mice were fed a control diet (CHOW) or a high-fat diet (HFD) for 15 weeks. During the final three weeks, FB1 (10 mg/kg bw/day) was added or not to the drinking water. mRNA expression levels of genes that encode for specific biological processes were analyzed in liver samples (n=12/group). (A) mRNA expression levels of genes representative of Lipogenesis process were studied through *Acca*; *Acly*; *Elovl6*; *Fasn*; *Gck*; *Lpk*; *Scd1* and *Srebp1-c* . (B) Typical mRNA expression levels of genes representative of inflammatory process were studied through *Tlr4*; *Myd88*; *Nlrp3*; *Pycard* and *Casp1*. (C) Illustrative mRNA expression levels of genes representative of fibrosis and NASH were respectively investigated through *Col3a1*; *Krt18* and *Panx1*. Results are presented as the mean ± SEM. #diet effect, *treatment effect; * or # p-value< 0.05, ** or ## p-value<0.01, *** or ### p-value<0.001; FB1: Fumonisin B1; CTRL: not exposed to FB1

**Figure S3. Effect of FB1 on sphingolipid homeostasis.**

C57BL/6J male mice were fed a control diet (CHOW) or a high-fat diet (HFD) for 15 weeks. During the final three weeks, FB1 (10 mg/kg bw/day) was added or not to the drinking water. (A) Relative expression of the most modulated gene associated to sphingolipid metabolism, plotted as the log2-fold change to their expression in the CHOW-CTRL group. Data were extracted from the liver gene expression analysis realized with Agilent microarrays. (B) Comparative hepatic level content of the sphingoid bases: Sphinganine and Sphingosine, and (C) their phosphate derivatives: Dihydro-sphingosine-1-phosphate and Sphingosine-1-phosphate. (D) Gangliosides total hepatic contents of: Glucosyl-ceramides; Lactosyl-ceramides; GM3 and GB3 and (E) their respective distribution according the length of their fatty acid chain residue. Results are the mean ± SEM (n=12/group except for Dihydro-sphingosine-1-phosphate for wich each level correspond to the pooling of 6 mouse samples) #diet effect, *treatment effect; * or # p-value< 0.05, ** or ## p-value<0.01, *** or ### p-value<0.001; FB1: Fumonisin B1; CTRL: not exposed to FB1; ND: Not Detected
