## Supplementary figures and images for "Obesity promotes Fumonisin B1 toxicity and induces hepatitis"

### Fig. S1

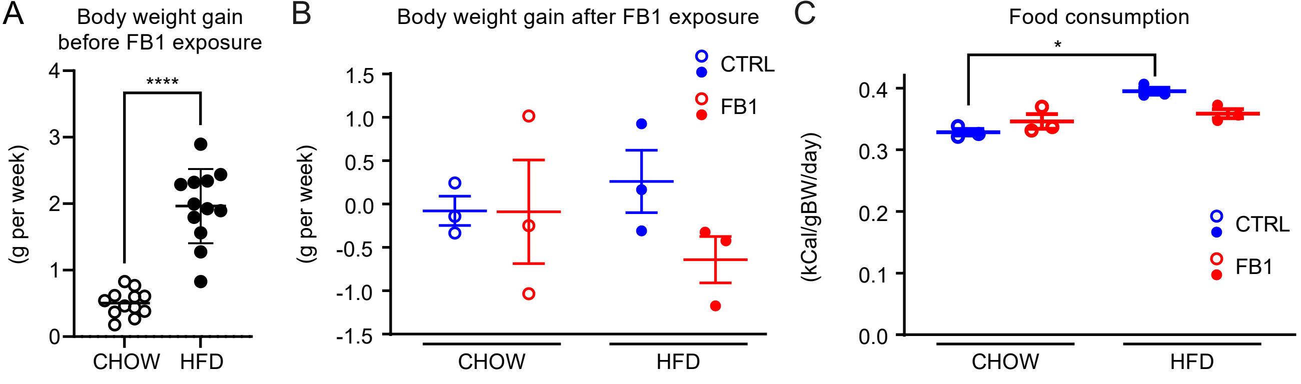

### Fig. S2

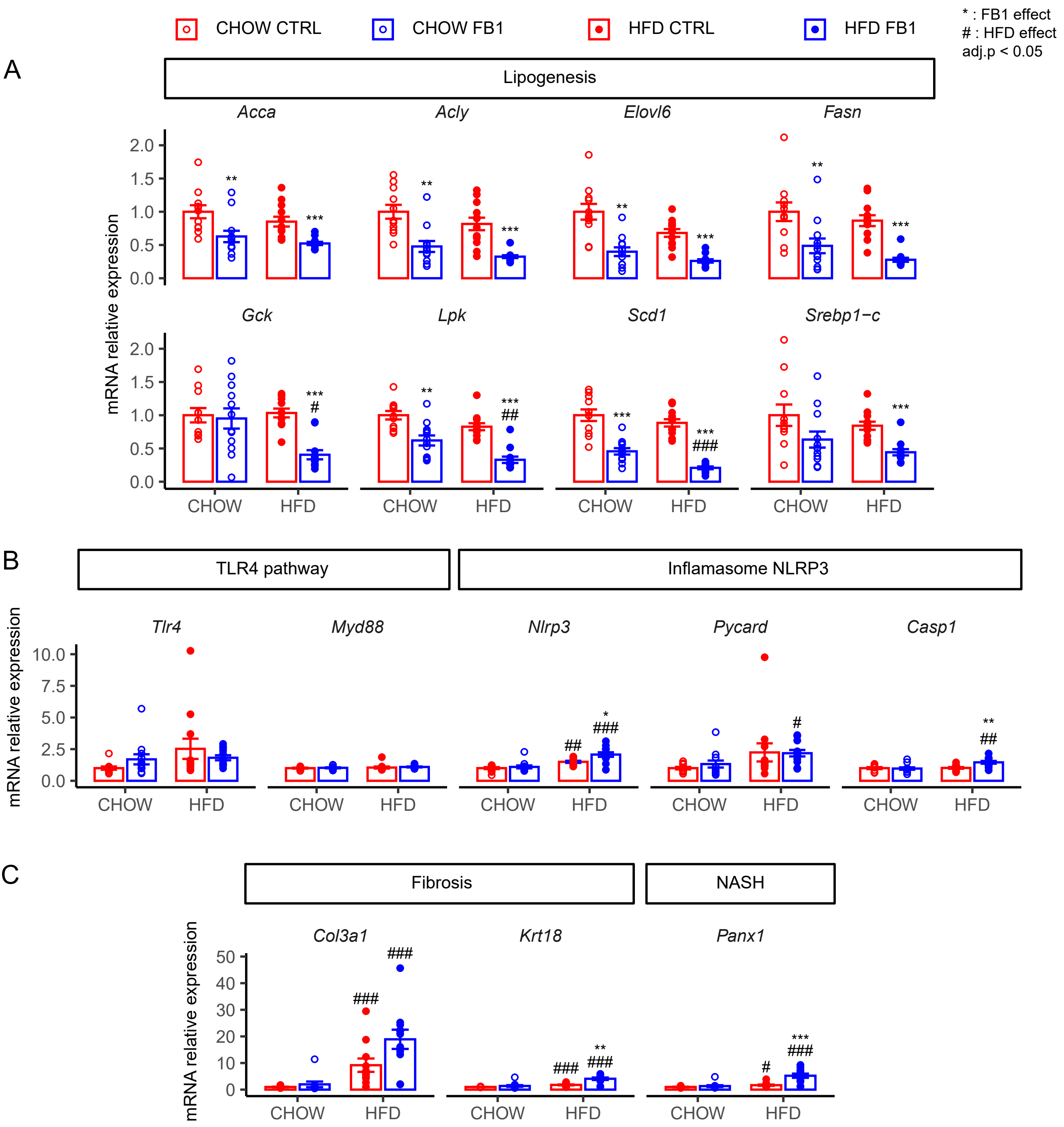

### Fig. S3

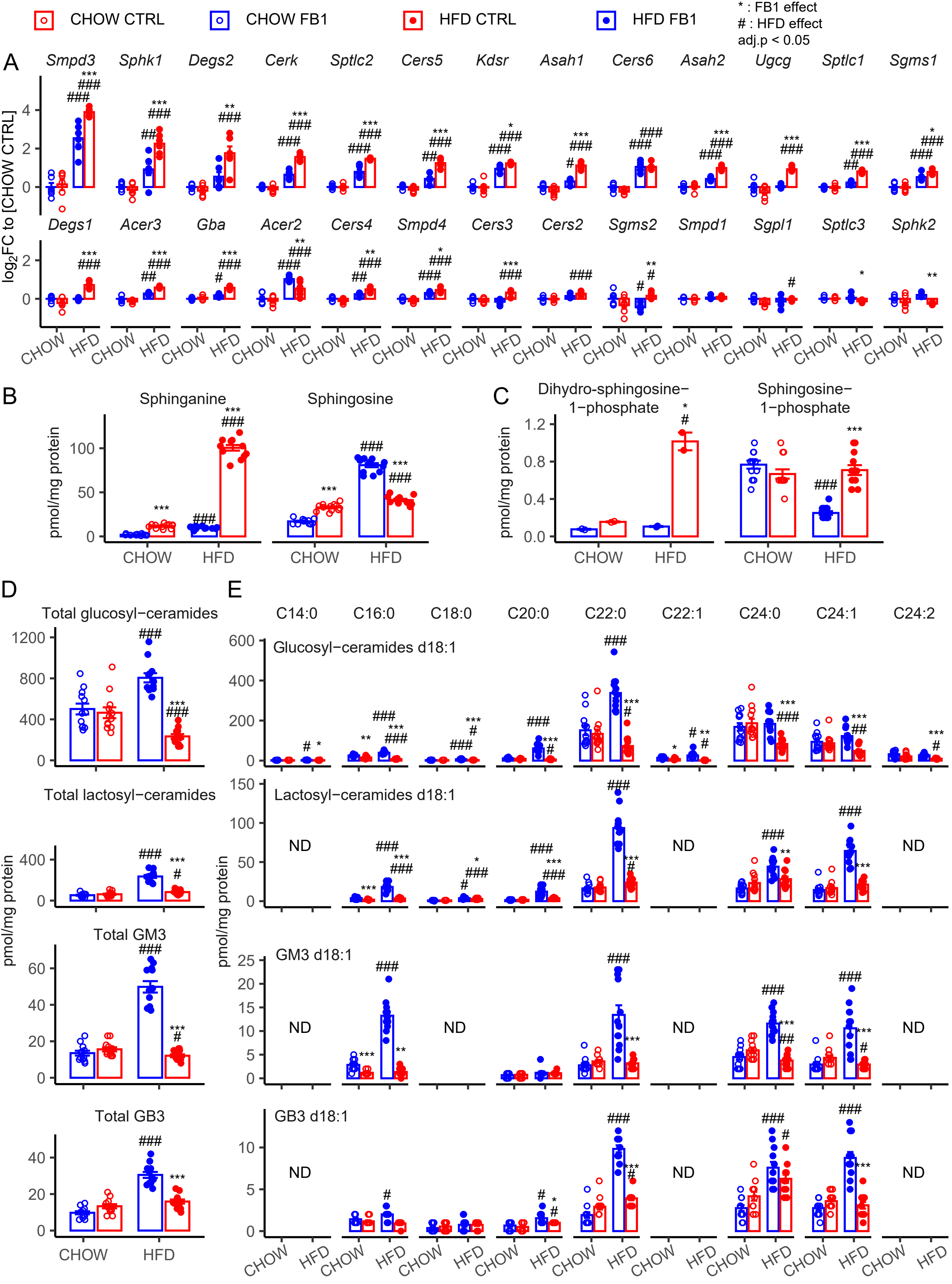
